# Functional and antigenic characterization of SARS-CoV-2 spike fusion peptide by deep mutational scanning

**DOI:** 10.1101/2023.11.28.569051

**Authors:** Ruipeng Lei, Enya Qing, Abby Odle, Meng Yuan, Timothy J.C. Tan, Natalie So, Wenhao O. Ouyang, Ian A. Wilson, Tom Gallagher, Stanley Perlman, Nicholas C. Wu, Lok-Yin Roy Wong

## Abstract

The fusion peptide of SARS-CoV-2 spike protein is functionally important for membrane fusion during virus entry and is part of a broadly neutralizing epitope. However, sequence determinants at the fusion peptide and its adjacent regions for pathogenicity and antigenicity remain elusive. In this study, we performed a series of deep mutational scanning (DMS) experiments on an S2 region spanning the fusion peptide of authentic SARS-CoV-2 in different cell lines and in the presence of broadly neutralizing antibodies. We identified mutations at residue 813 of the spike protein that reduced TMPRSS2-mediated entry with decreased virulence. In addition, we showed that an F823Y mutation, present in bat betacoronavirus HKU9 spike protein, confers resistance to broadly neutralizing antibodies. Our findings provide mechanistic insights into SARS-CoV-2 pathogenicity and also highlight a potential challenge in developing broadly protective S2-based coronavirus vaccines.

## INTRODUCTION

While the world is slowly returning to normal from the COVID-19 pandemic, severe acute respiratory syndrome coronavirus 2 (SARS-CoV-2) continues to circulate in the human population. Due to the importance in COVID-19 vaccine development, spike (S) is the most studied SARS-CoV-2 protein. S facilitates virus entry by binding to the host receptor angiotensin-converting enzyme 2 (ACE2) and mediates virus-host membrane fusion by undergoing drastic conformational changes^1^. Membrane fusion is activated by the cleavage of the S2’ site in the S2 domain by either TMPRSS2 at the cell surface or cathepsins in the endosome^2–4^. With cleavage of the S2’ site, the fusion peptide is exposed and inserted into the membrane of the host cell^5^. Subsequently, the S2 domain rearranges into a stable six-helix bundle with a long central three-stranded coiled coil to complete membrane fusion^6,7^. Although early SARS-CoV-2 variants enter cells mainly by TMPRSS2-mediated cleavage, some Omicron variants have been shown to utilize cathepsin-mediated endosomal entry^8–11^. This shift of cell entry pathway may associate with changes in cellular tropism and reduction in virulence^8,9^. As a result, studying the determinants of SARS-CoV-2 membrane fusion has important public health implications.

Residues 816-834 of the S protein, which locate immediately downstream of the S2’ cleavage site at Arg815/Ser816^12^, have generally been recognized as the bona fide SARS-CoV-2 fusion peptide (bFP, residues 816-834)^13–15^. Nevertheless, a recent cryo-EM structure of the postfusion SARS-CoV-2 S in a lipid bilayer membrane showed that the internal fusion peptide (iFP, residues 867-909) insert into the membrane, whereas the bFP was not resolved^16^. This observation appears to challenge the functional importance of bFP, but also indicates that additional analysis of the fusion peptide and fusion mechanism of SARS-CoV-2 S is warranted.

Neutralizing antibodies targeting the functionally important S2 domain have been isolated from convalescent individuals^17–21^. Unlike antibodies to the immunodominant receptor-binding domain (RBD) of S^22,23^, S2 antibodies typically have very broad cross-reactivity due to high S2 sequence conservation^17–21^. Neutralizing antibodies to an epitope that spans the S2’ cleavage site and the bFP can cross-react with diverse coronavirus strains from all four genera (α, β, γ and δ)^17,20,21,24^. These broadly neutralizing antibodies provide important insights into the development of a pan-coronavirus vaccine. However, comprehensive assessments of the genetic barrier for resistance to bFP antibodies have not been completed. Relatedly, the mutational tolerance of the SARS-CoV-2 bFP is largely elusive.

Deep mutational scanning, which combines saturation mutagenesis and next-generation sequencing, allows the phenotypes of many mutations to be measured in parallel. Deep mutational scanning has been applied to study the mutational fitness effects of various medically important RNA viruses, including influenza virus^25,26^, human immunodeficiency virus^27^, hepatitis C virus^28^, and Zika virus^29^. All of these viruses can be evaluated using efficient plasmid-based reverse genetic systems, which are pre-requisites for applying deep mutational scanning to study viral replication fitness. At the same time, most, if not all, deep mutational scanning studies of SARS-CoV-2 have been performed using protein display or pseudovirus systems^30–33^. Although these studies have offered critical insights into antibody resistance and biophysical constraints of SARS-CoV-2 evolution, they do not directly measure virus replication fitness or virulence. While multiple reverse genetic systems are available for SARS-CoV-2^34–36^, they are more complex than those for other RNA viruses, mainly due to the larger genome size of SARS-CoV-2. Thus, probing the fitness effects of SARS-CoV-2 mutations by deep mutational scanning can be technically challenging.

In this study, we performed deep mutational scanning of S residues 808-855, spanning the S2’ cleavage site, bFP, and fusion peptide proximal region (FPPR)^16^, using a bacterial artificial chromosome (BAC)-based reverse genetic system of SARS-CoV-2. Our results revealed that the bFP (residues 816-834) has a very low mutational tolerance. In addition, we identified mutations upstream of the S2′ cleavage site that reduced TMPRSS2-mediated entry. Further characterizations of these mutations suggested a relationship between sensitivity for TMPRSS2-mediated S2’ cleavage, cell entry pathway, and virus virulence. We also identified a mutation in the bFP that resists two broadly neutralizing bFP antibodies and naturally exists in a bat coronavirus strain.

## RESULTS

### Deep mutational scanning of SARS-CoV-2 bFP

Based on a BAC-based reverse genetic system of SARS-CoV-2 Wuhan-Hu-1 (pBAC SARS-CoV-2)^37,38^, we constructed a saturation mutagenesis library that contained all possible single amino acid mutations in the bFP and FPPR (residues 816-855) of the SARS-CoV-2 S, as well as the eight residues immediately upstream of the S2’ cleavage site (residues 808-815)^38^. The BAC mutant library was transfected into Vero cells to generate a virus mutant library, which was then passaged once in Calu-3 or Vero cells for 48 hours. The frequencies of individual mutations in the BAC mutant library and the post-passaged mutant library were determined by next-generation sequencing. The fitness value of each mutation was calculated based on its frequency enrichment and normalized such that the mean fitness values of silent mutations and nonsense mutations were 1 and 0, respectively (**see Methods**). The fitness values of 893 (98%) out of 912 all possible amino acid mutations across the 48 residues of interest were measured (**Figure 1**). Pearson correlation coefficients of 0.62 (Calu-3) and 0.58 (Vero) were obtained between two biological replicates (**Figure S1A-B**), demonstrating the reproducibility of our deep mutational scanning experiments. Moreover, the fitness value distributions of silent and nonsense mutations had minimal overlap, further validating our results.

**Figure 1.**
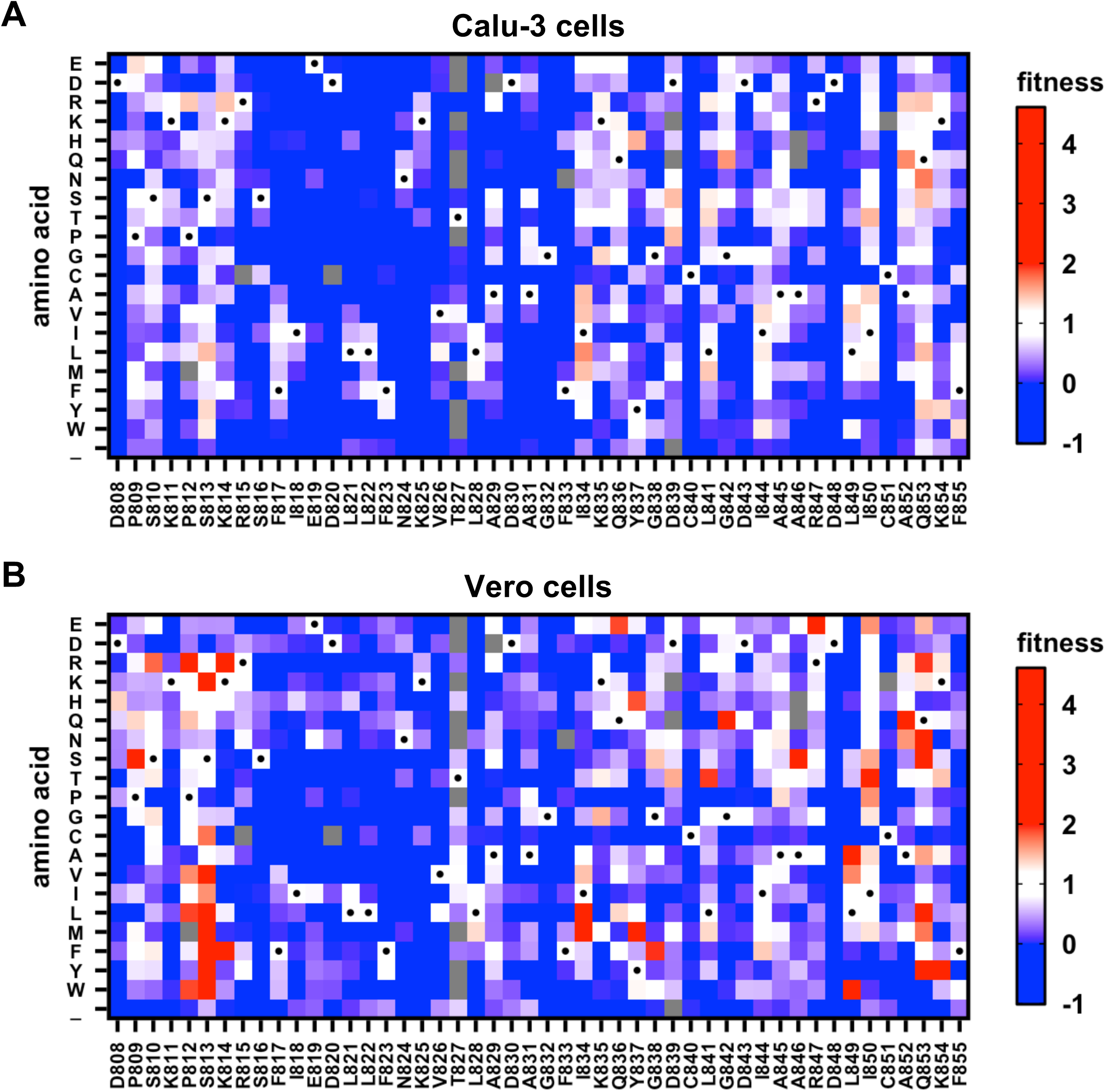
Deep mutational scanning of SARS-CoV-2 bFP and FPPR. The fitness values of individual mutations at residues 808 to 855 of SARS-CoV-2 S were measured by deep mutational scanning in **(A)** Calu-3 cells and **(B)** Vero cells and are shown as heatmaps. Wild-type (WT) amino acids are indicated by black circles. “_” indicates nonsense mutations. Mutations in gray were excluded in our data analysis due to low frequency in the plasmid mutant library. Red indicates superior fitness, white similar to WT, and blue reduced fitness.

Recently, the effects of ∼7,000 natural mutations of SARS-CoV-2 S on cell entry have been quantified by a pseudovirus-based deep mutational scanning experiment^33^. The fitness effects of natural mutations in the SARS-CoV-2 genome have also been estimated using a phylogenetic-based approach in another study^39^. Although these studies only examined <50% of all possible amino acid mutations from residues 808 to 855, their measurements moderately correlated with our deep mutational scanning results (rank correlation ranges from 0.36 to 0.49, **Figure S1G-J**).

### Mutations at residue 813 modulate protease utilization for S2’ cleavage

Based on our deep mutational scanning results, we observed that certain mutations had high fitness values in Vero cells but not in Calu-3 cells (**Figure 1**). This observation was particularly apparent at residue 813, which is upstream of the S2′ cleavage site. Coronaviruses, including SARS-CoV-2, are known to enter Calu-3 cells through TMPRSS2-mediated membrane fusion on the cell surface^10,40,41^. In contrast, coronaviruses enter Vero cells, with low TMPRSS2 expression, through cathepsin-mediated membrane fusion in endosomes^10,41–43^. As a result, we hypothesized that mutations at residue 813 shifted the preference of protease utilization for the S2’ cleavage site.

To test this hypothesis, we generated VSV-based pseudoparticles (VSVpps) bearing wild-type (WT), S813V, or S813K SARS-CoV-2 S. Although S813V and S813K slightly decreased the incorporation of S into VSVpp (**Figure S2A**), their efficiency of Vero cell entry was similar to WT (**Figure 2A**). However, when Vero cells overexpressed TMPRSS2 (Vero-TMPRSS2), both S813V and S813K had reduced entry compared to WT (**Figure 2B**), suggesting that mutations at residue 813 decreased sensitivity to TMPRSS2-mediated activation. Furthermore, cathepsin inhibitor E64D, but not TMPRSS2 inhibitor camostat, significantly reduced Vero cell entry to a greater extent in S813V and S813K compared to WT (**Figure 2C-D**). In contrast, camostat reduced Vero-TMPRSS2 cell entry to similar extents among WT, S813V, and S813K (**Figure S2C**), indicating that TMPRSS2-mediated entry was preferred when TMPRSS2 was overexpressed. This same experiment was then performed in the presence of fetal bovine serum (FBS), which suppresses cell surface protease-mediated (e.g. TMPRSS2-mediated) entry^44^ (**Figure S2B**). When FBS was added, Vero-TMPRSS2 cell entry of S813V and S813K became less sensitive to camostat, and hence with less reliance on TMPRSS2, compared to WT (**Figure S2D**). As a control, we also demonstrated that Calu-3 cell entry was camostat-sensitive and hence TMPRSS2-dependent (**Figure S2E**), which agrees with previous studies^10,40,41^. Taken together, these results suggest that S813V and S813K have reduced TMPRSS2-mediated entry.

**Figure 2.**
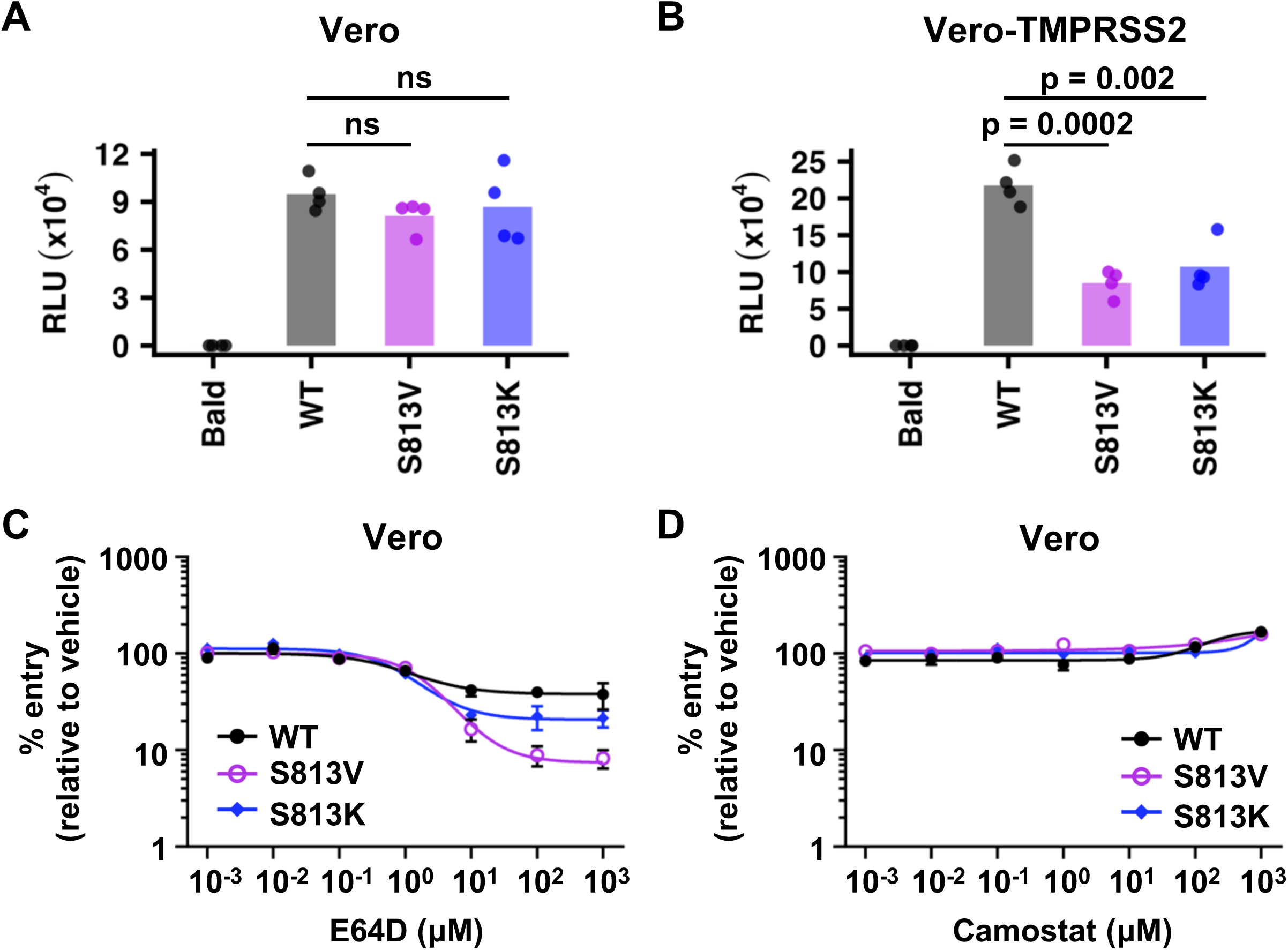
Mutations at residue 813 influence the protease utilization during cell entry. **(A)** Vero cell entry of VSVpps bearing various SARS-CoV-2 S constructs was measured by the relative light unit (RLU) in a luciferase assay. **(B)** Vero-TMPRSS2 cell entry of VSVpps bearing various SARS-CoV-2 S constructs. Each bar represents the mean of four biological replicates. Each datapoint represents one biological replicate. Deviations from the WT were analyzed by two-sample t-tests. “ns” indicates not significance (i.e. p-value > 0.05). **(C and D)** The effects of **(C)** E64D (cathepsin inhibitor) or **(D)** camostat (TMPRSS2 inhibitor) on Vero cell entry of VSVpps bearing various SARS-CoV-2 S constructs are shown. Curves depicted in (**C**), are significantly different (p = 0.0088, two-way ANOVA). Mean and standard error of the mean (SEM) of four independent biological replicates are depicted.

While S813V and S813K mutants entered Vero cells as efficiently as VSVpps with WT S proteins (**Figure 2A**), they had higher fitness values than WT in the deep mutational scanning experiment (**Figure 1B**). This seeming discrepancy may be explained by differences between the experimental systems. The deep mutational scanning was based on the recombinant SARS-CoV-2, whereas the VSVpp experiment only measured the efficiency of cell entry, which did not represent the entire virus life cycle. Besides, the incorporation efficiency and density of S on the virion were likely different between VSVpp and recombinant SARS-CoV-2. Despite these differences, both the VSVpp and deep mutational scanning experiments support the hypothesis that mutations at residue 813 modulate the sensitivity to TMPRSS2-mediated activation of virus entry.

### Mutations at residue 813 decrease SARS-CoV-2 virulence *in vivo*

To investigate the effects of S813V and S813K in authentic SARS-CoV-2, we introduced the two mutations individually into a mouse-adapted SARS-CoV-2 strain^38^. Vero cells, Vero-TMPRSS2 cells, and Vero cells overexpressing both TMPRSS2 and ACE2 (Vero-TMPRSS2/ACE2) were simultaneously infected with the same aliquot of virus. The numbers of plaques obtained for WT, S813V, and S813K mutants were all enhanced in Vero-TMPRSS2 and Vero-TMPRSS2/ACE2 cells as compared to Vero cells. However, such enhancement was significantly higher for WT than S813V and S813K mutants in both Vero-TMPRSS2 and Vero-TMPRSS2/ACE2 cells (**Figure 3A-B**). This observation substantiates the conclusion that that S813V and S813K exhibit reduced sensitivity to TMPRSS2-mediated cleavage. Consistently, the S813V mutant also showed significantly higher titer than WT at 24 hours post-infection (hpi) in Vero cells (p = 0.01, **Figure 3C**), but not in Vero-TMPRSS2 cells (**Figure 3D**).

**Figure 3.**
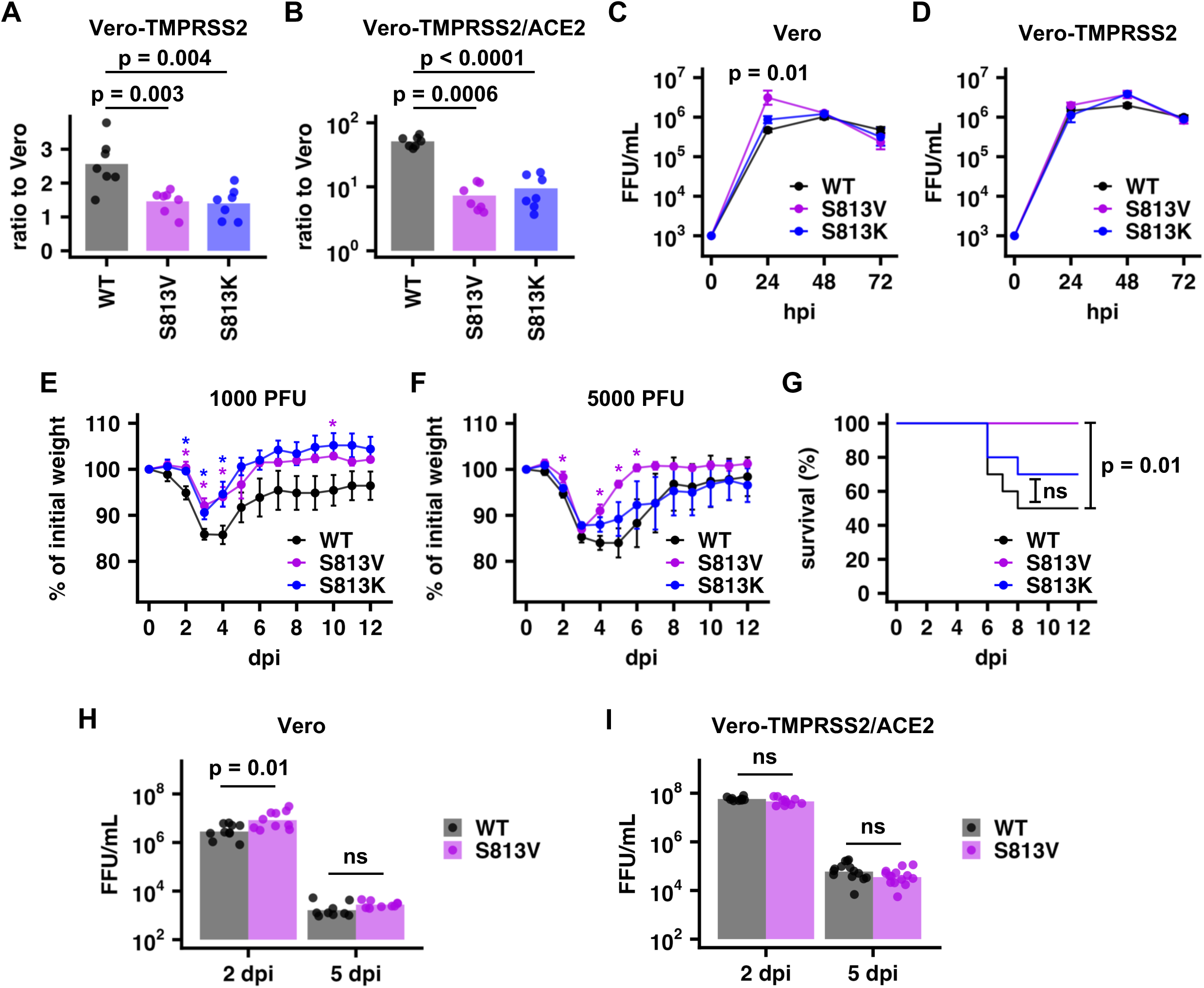
S813V mutation reduces virulence *in vivo*. **(A-B)** Vero, Vero-TMPRSS2, and Vero-TMPRSS2/ACE2 cells were separately infected with WT, S813V or S813K viruses from the same aliquot for each virus. The numbers of plaques obtained from **(A)** Vero-TMPRSS2 cells or **(B)** Vero-TMPRSS2/ACE2 cells were normalized to those obtained from Vero cells. Bar represents the mean of seven biological replicates. Each datapoint represents one biological replicate. P-values were computed by two-sample t-tests. **(C)** Vero cells or **(D)** Vero-TMPRSS2 cells were infected with WT, S813V, or S813K mutants at a multiplicity of infection of 0.01. Virus titers were determined for each variant at the indicated time point. Each data point represents the geometric mean of three biological replicates and the error bar represents geometric standard deviation (SD). Representative data from two independent experiments are shown. Deviations from the WT were analyzed by two-sample t-tests. **(E-F)** Percentage of initial weight change of C57BL/6 mice (n = 5 to 10) infected with **(E)** 1000 PFU or **(F)** 5000 PFU of WT, S813V, or S813K mutants. Data points in weight curve represent the mean and error bars represent the SEM. Deviations from the WT were analyzed by two-sample t-tests. “*” indicates p-value < 0.01. **(G)** Kaplan-Meier survival curves are shown for C57BL/6 mice infected with 5000 PFU of S813V, or S813K mutants. “ns” indicates not significant (i.e. p-value > 0.05). Of note, all mice infected with 1000 PFU of WT, S813V, or S813K mutants survived. **(H-I)** Virus titers in the lungs of mice infected with 5000 PFU of WT, S813V, or S813K mutants were measured at the indicated time point on **(H)** Vero cells and **(I)** Vero-TMPRSS2/ACE2 cells. Bars represent the geometric mean and error bars represent the geometric SD. dpi, days post-infection.

We next aimed to understand the effects of mutations at residue 813 on virulence in mice. C57BL/6 mice were infected with 1000 or 5000 plaque-forming units (PFU) of WT, S813V, or S813K mutants. At 1000 PFU, infection with either the S813V or S813K mutant caused significantly less weight loss compared to WT (**Figure 3E**). At 5000 PFU, the S813V mutant virus again caused less weight loss than WT (**Figure 3F-G**), despite having similar, if not higher virus titers in the lungs at 2 and 5 days post-infection (dpi) compared to WT (**Figure 3H-I**). Together, these data indicate that mutations at residue 813 decreased virulence *in vivo*.

### Low mutational tolerance of the bFP

Although some mutations, such as those at residue 813, showed differential fitness effects between Calu-3 and Vero cells, many mutations in the deep mutational scanning experiment had consistently low fitness values between the two cell lines (**Figure 1**). Subsequently, we aimed to identify regions with low mutational tolerance. Here, we defined the mutational tolerance at each residue as the average fitness value of mutations at the given residue in Calu-3 cells. Residues that interact with the host membrane should have lower mutational tolerance due to functional constraints, as demonstrated by a previous deep mutational scanning study on influenza hemagglutinin (**Figure S3**)^45,46^. Notably, residues 816 to 833, which spanned most of the bFP, had low mutational tolerance (**Figure 4A**). In contrast, the FPPR had a much higher mutational tolerance.

**Figure 4.**
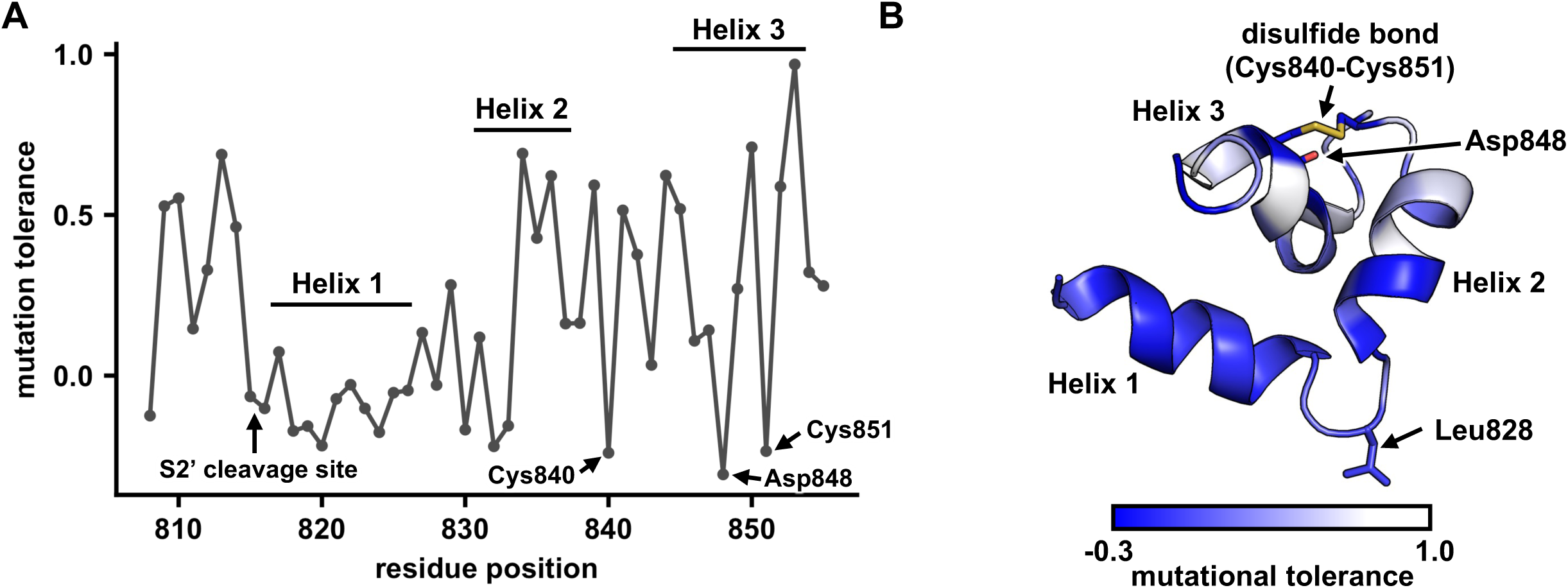
Structural analysis of the mutational tolerance of SARS-CoV-2 bFP and FPPR. **(A)** Mutational tolerance of each residue in Calu-3 cells is shown on the NMR structure of the bFP and FPPR (PDB 7MY8)^13^. A disulfide bond (yellow in panel B) is present in the FPPR between Cys840 and Cys851. **(B)** The mutational tolerance of each residue in Calu-3 cells is shown. The locations of helices 1-3 in the NMR structure of the bFP and FPPR (PDB 7MY8)^13^ are indicated. The side chains of Leu828, Cys840, Asp848, and Cys851 are shown in stick representation.

An NMR structure of the bFP and FPPR indicates that they form a three-helix wedge-shaped structure when interacting with the host membrane, with Leu828, which locates between helix 1 and helix 2, pointing towards the interior of the membrane^13^. Based on the mutational tolerance data, we further propose that helix 1 and the N-terminal half of helix 2, which represent the bFP, could interact with the membrane during virus-host membrane fusion. In contrast, the C-terminal of helix 2 and helix 3, which represent the FPPR, would likely remain in the aqueous phase (**Figure 4B**). As a result, our deep mutational scanning data substantiates that the bFP interacts with the host membrane^13,47^.

We also identified three residues in the FPPR that had low mutational tolerance, namely Cys840, Asp848, and Cys851 (**Figure 4A**). The low mutational tolerance of Cys840 and Cys851 could be explained by the disulfide bond between them (**Figure 4B**). On the other hand, the functional importance of Asp848 was not as clear. Previous studies suggest that the bFP and FPPR each bind to a calcium ion via their negatively charged residues to promote membrane fusion^13,48^. All three negatively charged residues in the bFP, namely E819, D820, and D830, had very low mutational tolerance, consistent with these three residues representing the calcium-binding site in the bFP^48^. Our mutational tolerance data further suggested that Asp848 was the calcium-binding site in the FPPR (**Figure 4A**), since it was the only negatively charged residue in the FPPR that could not tolerate any non-negatively charged mutations (**Figure 1**). Consistently, Asp848, but not Asp839 and Asp843, which are the other two negatively charged residues in the FPPR, is conserved across all four genera of coronaviruses^49^.

### Resistance of F823Y mutation to bFP antibodies

Previous studies have shown that antibody resistance mutations can be identified by deep mutational scanning^26,50,51^. To investigate whether SARS-CoV-2 bFP can acquire resistance mutations to bFP antibodies, deep mutational scanning was performed in the presence of bFP antibodies COV44-62 and COV44-79, both of which can neutralize SARS-CoV-2 and cross-react with coronavirus strains from different genera^17^. These two antibodies engage the bFP differently and are encoded by different germline genes^17^. COV44-62 is encoded by IGHV1-2/IGLV2-8, whereas COV44-79 is encoded by IGHV3-30/IGKV1-12^17^.

Our deep mutational scanning results indicated that F823Y, which had minimal fitness cost (**Figure 1**), was a resistance mutation to both COV44-62 and COV44-79 (**Figure 5A-B, Figure S1C-F and Figure S4A**). To validate this finding, we generated VSVpp bearing SARS-CoV-2 S with the F823Y mutation. F823Y did not affect incorporation of SARS-CoV-2 S into VSVpp, S1-S2 stability, or cleavage at the S1/S2 site (**Figure S2A**). Nevertheless, F823Y S-bearing VSVpp conferred resistance to both COV44-62 and COV44-79 in a neutralization assay (**Figure 5C-D**). The resistance of F823Y appeared to be stronger against COV44-79 than COV44-62, since F823Y S-bearing VSVpp was partly neutralized by COV44-62, but not COV44-79 at the highest tested concentration (500 μg/mL) (**Figure 5C-D**). Consistently, F823Y mutation weakened the binding of an epitope-containing peptide to COV44-62 and COV44-79 by 8-fold and >40-fold, respectively (**Figure S4B**). These results demonstrate that resistance to bFP antibodies can be conferred by a single mutation.

**Figure 5.**
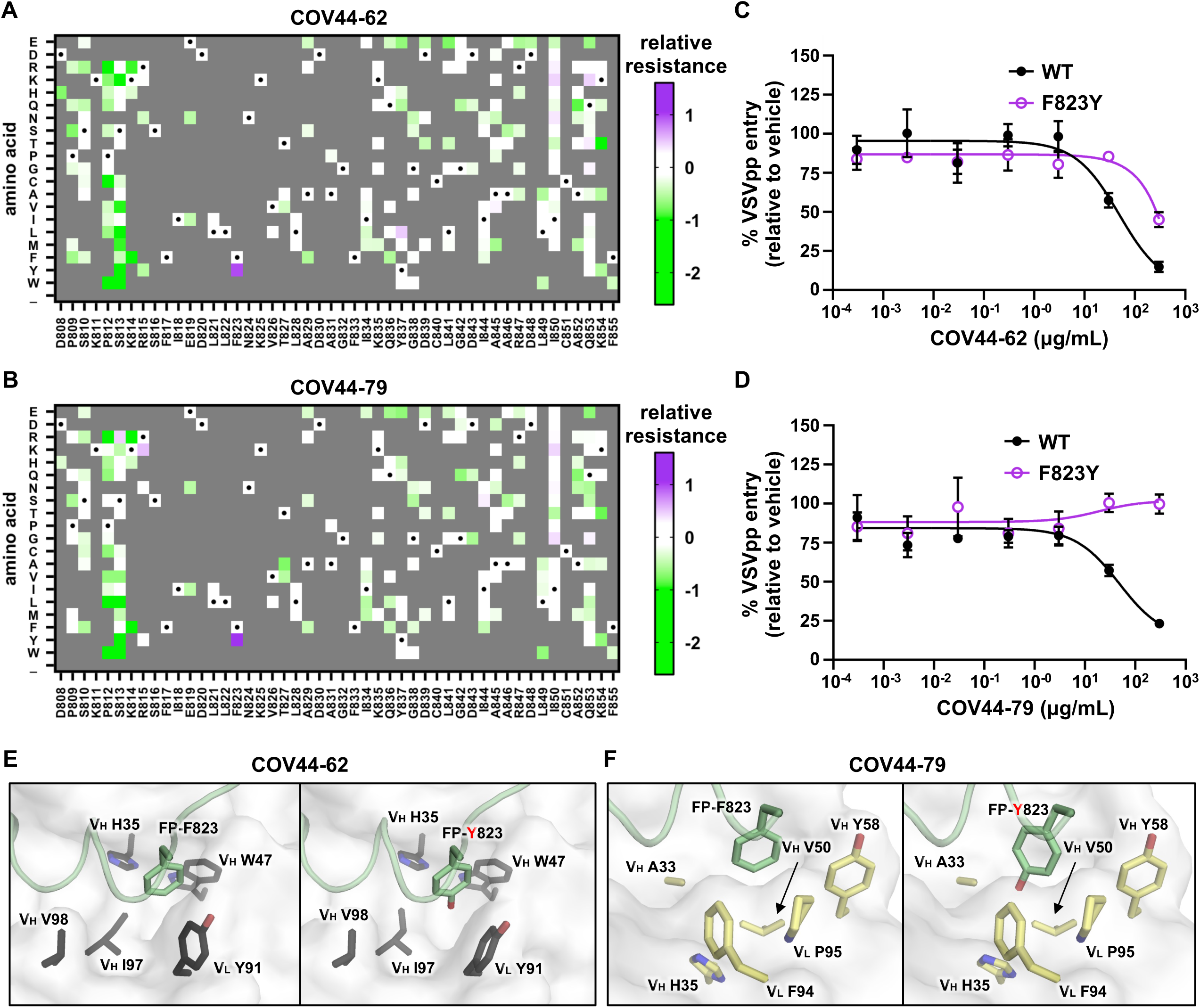
F823Y weakens binding of bFP antibodies. **(A-B)** Relative resistance for each mutation against **(A)** 230 μg/mL COV44-62 or **(B)** 330 μg/mL COV44-79 in Vero cells is shown as heatmaps. Relative resistance for WT is set as 0. Mutations with a fitness value of less than 0.75 in the absence of antibody are shown as gray. Amino acids corresponding to the WT sequence are indicated by the black dots. “_” indicates nonsense mutations. **(C-D)** The neutralization activities of **(C)** COV44-62 and **(D)** COV44-79 against VSVpp bearing WT or F823Y S are shown. Mean and SEM of three biological replicates are depicted. **(E-F)** The structural effects of F823Y on the binding of **(E)** COV44-62 (PDB 8D36)^17^ and **(F)** COV44-79 (PDB 8DAO)^17^ were modelled using FoldX^52^.

To understand the structural mechanism of antibody resistance, we further analyzed the previously determined x-ray crystal structures of COV44-62 and COV44-79 in complex with SARS-CoV-2 bFP^17^. FoldX was used to model the structural effect of F823Y mutation^52^. The major difference between Phe and Tyr is an extra side-chain hydroxyl group on Tyr. Our models showed that the hydroxyl group of Tyr823 pointed towards the bottom of hydrophobic pockets formed in the COV44-62 and COV44-79 binding sites (**Figure 4E-F**). Burying a polar hydroxyl group on Tyr side chain without forming any H-bond would impose an appreciable desolvation energy cost^53^. Consistently, FoldX indicated that F823Y mutation weakened the binding energy of COV44-62 and COV44-79 by 1.0 kcal/mol and 1.2 kcal/mol, respectively. These observations provide a mechanistic basis of the resistance to bFP antibodies conferred by F823Y.

## DISCUSSION

Most studies of the SARS-CoV-2 S protein focus on the RBD, since it is immunodominant and engages the host receptor ACE2 for cell entry^1,22,23^. In contrast, the S2 domain is less well characterized. Our study here provides important insights into how mutations in the regions adjacent to the S2’ cleavage site can modulate the preference of cell entry pathway as well as promote resistance to broadly neutralizing antibodies. Our results also advance the knowledge on the evolutionary potential of the SARS-CoV-2 S2 domain and demonstrate the feasibility of applying deep mutational scanning to authentic SARS-CoV-2.

A key result in this study is the low mutational tolerance of the bFP, which substantiates its functional importance during membrane fusion^13–15^. However, the bFP was located outside the membrane as a disordered region in a recent cryo-EM structure of postfusion SARS-CoV-2 S in a lipid bilayer membrane^16^. Instead, the iFP inserts into the membrane in this cryo-EM structure. However, the postfusion SARS-CoV-2 S in this cryo-EM structure does not have a cleaved S2’ site that is essential for membrane fusion during virus entry^16,54^. Besides, this cryo-EM structure was determined at pH 7.5 without any calcium ions, while SARS-CoV-2 S-mediated membrane fusion requires an acidic pH^55^ and the presence of calcium ions^48,56^. While it is always challenging for structural and biophysical studies of viral fusion proteins to emulate the physiological states as would occur *in vivo*, it is possible that both bFP and iFP of SARS-CoV-2 S interact with the host membrane^13,16^, but at different stages of the membrane fusion process. Future studies are therefore needed to better characterize the molecular mechanisms of the highly dynamic S-mediated membrane fusion process.

Another major observation in our study is that mutations at SARS-CoV-2 S residue 813 influenced host cell entry and sensitivity to TMPRSS2-mediated S2’ cleavage. Consistently, similar findings on residue 813 have recently been described for different SARS-CoV-2 variants as well as SARS-CoV^57^. Previous studies showed that H655Y and N969K mutations in Omicron can shift the preference from TMPRSS2-mediated cell surface entry to cathepsins-mediated endosomal entry, resulting in reduced virulence^8–10^. The proposed underlying mechanism is that they stabilize the S trimer, and hence decrease the fusogenicity and cell surface entry efficiency^9–11^. We also observed this relationship between cell entry pathway and virulence in mutations at residue 813. However, unlike H655Y and N969K, residue 813 is near the S2’ cleavage site (**Figure S5**). Therefore, while S813V and S813K have similar phenotypes as H655Y and N969K, their molecular mechanisms are unlikely to be the same. Given that other residues flanking the S2’ cleavage site can also modulate protease preference for S2’ cleavage^58,59^, mutations in this region may provide valuable information on the preference of cell entry pathway and pathogenicity as SARS-CoV-2 continues to evolve.

There are currently five coronavirus strains circulating in the human population (229E, HKU1, NL63, OC43, and SARS-CoV-2). In addition, other zoonotic coronaviruses continue to pose a pandemic threat^60^. As a result, developing a pan-coronavirus vaccine has become an attractive idea, especially after the discovery of broadly neutralizing antibodies to the bFP^17,20,21,24^. However, despite the high sequence conservation of the bFP, our study here found that F823Y mutation can confer strong resistance against bFP antibodies. F823Y is a natural variant in bat betacoronavirus HKU9 and is also observed in circulating SARS-CoV-2 at a very low frequency (**Figure S6**). Although these observations represent a potential obstacle for the development of a pan-coronavirus vaccine, resistance mutations against bFP antibodies are rare in our deep mutational scanning results, partly due to the high fitness cost of most mutations in the bFP. Therefore, we concur that the bFP is a promising target for the development of a pan-coronavirus vaccine^17,20,21,24^.

## ACKNOWLEDGEMENTS

This work was supported by National Institutes of Health (NIH) R21 AI 178391 (T.G.), K99 AI170996 (L.-Y.R.W.), P01 AI060699 (S.P.), R01 AI129269 (S.P.), the Searle Scholars Program (N.C.W.) and the Bill and Melinda Gates Foundation grant INV-004923 (I.A.W.).

## AUTHOR CONTRIBUTIONS

L.-Y.R.W., R.L., E.Q., T.G., S.P., and N.C.W. conceived and designed the study. L.-Y.R.W., R.L., A.O. performed the deep mutational scanning experiments. N.C.W. and N.T.Y.S analyzed the deep mutational scanning data. R.L. and T.J.C.T. expressed and purified the antibodies. E.Q. performed the functional characterization experiments. M.Y. and I.A.W. performed the biolayer interferometry experiment. L.-Y.R.W. and N.C.W. wrote the paper and all authors reviewed and/or edited the paper.

## METHODS

### Cell lines

HEK293T, Vero, Vero-TMPRSS2 and Vero-TMPRSS2/ACE2 cells were maintained in Dulbecco’s Modified Eagle Media (DMEM) containing 10 mM HEPES, 100 nM sodium pyruvate,

0.1 mM non-essential amino acids, 100 U/ml penicillin G, and 100 µg/ml streptomycin, and supplemented with 10% fetal bovine serum (FBS, Atlanta Biologicals). Calu-3 cells were maintained in Minimum Essential Media (MEM) supplemented with 20% FBS, 100 U/ml penicillin G, and 100 µg/ml streptomycin. All cell lines were cultured in a 5% CO_2_ incubator at 37°C.

### SARS-CoV-2 infection of mice

C57BL/6 mice of both sexes at 4 to 6 months old were used in this study. Mice were anaesthetized with ketamine-xylazine and infected intranasally with the indicated amount of virus in a total volume of 50 μL DMEM. Animal weight and health were monitored daily. All experiments with SARS-CoV-2 were performed in a biosafety level 3 (BSL3) laboratory at the University of Iowa. All animal studies were approved by the University of Iowa Animal Care and Use Committee and meet stipulations of the Guide for the Care and Use of Laboratory Animals.

### Virus titer by plaque assay

At the indicated times, mice were euthanized and transcardially perfused with PBS. Lungs were collected and homogenized before clarification by centrifugation and tittering. Virus or tissue homogenate supernatants were serially diluted in DMEM. Vero, Vero-TMPRSS2 or Vero-TMPRSS2/ACE2 cells in 12-well plates were inoculated at 37°C in 5% CO_2_ for 1 h and gently rocked every 15 min. After removing the inocula, plates were overlaid with 0.6% agarose containing 2%LFBS. After 3 days, overlays were removed, and plaques visualized by staining with 0.1% crystal violet. Viral titers were quantified as PFULper mL tissue.

### Virus titer by focus forming assay

Virus or tissue homogenate supernatants were serially diluted in DMEM. Vero, Vero-TMPRSS2 or Vero-TMPRSS2/ACE2 cells in 96-well plates were inoculated at 37°C in 5% CO_2_ for 1Lh and gently rocked every 15 min. After removing the inocula, plates were overlaid with 1.2% methylcellulose containing 2%LFBS. The next day, overlays were removed, and cells stained with anti-nucleocapsid antibody for SARS-CoV-2 for 1 h at 37°C and then with HPR-conjugated secondary antibody for 1 h at 37°C. Foci were visualized by peroxidase substrate. Viral titers were quantified as fluorescent focus unit (FFU)Lper mL tissue.

### Virus growth assay

Vero or Vero-TMPRSS2 cells in 12-well plates were infected with 0.01 MOI of the indicated virus diluted in DMEM. Cells were frozen at the indicated time points. Virus titers were determined by either plaque assay or focus forming assay. Three biological replicates were included for each time point.

### Mutant library construction

Mutant library of residues 808-855 of SARS-CoV-2 S was constructed based on a BAC-based reverse genetic system of SARS-CoV-2 Wuhan-Hu-1 (p-BAC SARS-CoV-2)^37,38^. Saturation mutagenesis was performed using an overlapping PCR strategy as described previously^32^. Briefly, a library of mutant inserts was generated by two separate batches of PCRs to cover the entire region of interest (residues 808-855). The first batch of PCRs consisted of 6 reactions, each containing one cassette of forward primers and the universal reverse primer 5’-GAC TGG AGT TCA GAC GTG TGC TCT TCC GAT CTT TGA GCA ATC ATT TCA TCT GTG AG-3’.

Each cassette contained an equal molar ratio of eight forward primers that had the same 21 nucleotides (nt) at the 5’ end and 15 nt at the 3’ end. Each primer within a cassette was also encoded with an NNK (N: A, C, G, T; K: G, T) sequence at a specified codon positions for saturation mutagenesis. In addition, each primer also carried unique silent mutations (also known as synonymous mutations) to help distinguish between sequencing errors and true mutations in downstream sequencing data analysis as described previously^61^. The forward primers, named as CassetteX_N (X: cassette number, N: primer number), are listed in **Table S1**. The second batch of PCR consisted of another 6 PCRs, each with a universal forward primer 5’-CAC TCT TTC CCT ACA CGA CGC TCT TCC GAT CTT TTG GTG GTT TTA ATT TTT CAC AA-3’ and a unique reverse primer as listed in **Table S1**. Subsequently, 6 overlapping PCRs were performed using the universal forward and reverse primers, as well as a mixture of 10 ng each of the corresponding products from the first and second batches of PCR. The 6 overlap PCR products were then mixed at equal molar ratio to generate the final insert of the mutant library. All PCRs were performed using PrimeSTAR Max polymerase (Takara Bio, catalog no. R045B) per the manufacturer’s instruction, followed by purification using the Monarch Gel Extraction Kit (New England Biolabs, catalog no. T1020L).

The FPPR mutant library PCR product was introduced into SARS-CoV-2 BAC encoding Wuhan-Hu-1 sequence by a two-step linear lambda red recombination process^62,63^. The first step removed and replaced the region of interest with GalK-Kan selection marker while the second step removed and replaced the GalK-Kan selection marker with the mutant library PCR product. In brief, GalK-Kan selection marker flanked by SARS-CoV-2 sequence was PCR-amplified from pYD-C225^62^ and gel-purified. Gel-purified GalK-Kan fragments were transformed into SW102 cells carrying the SARS-CoV-2 BAC by electroporation for linear lambda red recombination. Recombinants were selected by Kanamycin resistance culture plates. The presence of GalK-Kan cassette in selected recombinants was verified by PCR with primers flanking the area of recombination: 5’-CCA TAC CCA CAA ATT TTA CTA TTA GTG TTA CCA CA-3’ and 5’-TTG ACC ACA TCT TGA AGT TTT CCA AGT G-3’). Verified recombinants were further introduced with the mutant library PCR product by electroporation for a second round of linear lambda red recombination. Two electroporation was performed separately to obtain two independent BAC mutant libraries as replicates. Successful recombinants were selected using 2-deoxy-galactose-based culture plates. All viable clones were collected and pooled to generate the BAC mutant library. The loss of the GalK-Kan cassette (and hence the FPPR sequence) in the BAC mutant library was confirmed by PCR with primers flanking the area of recombination: 5’-CCA TAC CCA CAA ATT TTA CTA TTA GTG TTA CCA CA-3’ and 5’-TTG ACC ACA TCT TGA AGT TTT CCA AGT G-3’. GalK-Kan selection markers were amplified with primers: 5’-ATG TAC ATT TGT GGT GAT TCA ACT GAA TGC AGC AAT CTT TTG TTG CAA TA<u>C CTG TTG ACA ATT AAT CAT CG</u>-3’ and 5’-GCC AAT AGC ACT ATT AAA TTG GTT GGC AAT CAA TTT TTG GTT CTC ATA GA<u>C TCA GCA AAA GTT CGA TTT A</u>-3’. Sequences complementary to pYD-C225 are underlined.

S813V and S813K were first individually introduced to an expression construct encoding SARS-CoV-2 S with NEB Q5 site-directed mutagenesis kit. S813K was introduced with primers: 5’-ATC AAA ACC A<u>AA G</u>AA GAG GTC ATT TAT TG-3’ and 5’-GGA TCT GGT AAT ATT TGT G-3’; S813V was introduced with primers: 5’-ATC AAA ACC A<u>GT G</u>AA GAG GTC ATT TAT TGA AG-3’ and 5’-GGA TCT GGT AAT ATT TGT G-3’. The mutated codons for S813K and S813V are underlined. The part of the S protein encoding S813K or S813V were separately amplified with primers: 5’-CCA TAC CCA CAA ATT TTA CTA TTA GTG TTA CCA CA-3’ and 5’-TTG ACC ACA TCT TGA AGT TTT CCA AGT G-3’ from the expression construct of the SARS-CoV-2 S encoding S813K or S813V generated from the site-directed mutagenesis process. The PCR products were introduced into SARS-CoV-2 BAC as described above.

### Rescue and passage of the viral mutant library

2 μg of BAC mutant library were transfected into Vero cells with Lipofectamine 3000 (Thermo Fisher Scientific, catalog #: L3000008) into each well of a 6-well plate according to manufacturer’s protocol (12 μg in total for each replicate). Cells were monitored daily for cytopathic effects (CPE). Cultures were harvested when CPE was >50% by freezing at −80°C. Viruses rescued from each well of the transfected 6-well plate were pooled independently for each replicate to generate the P0 virus. The titers for P0 virus were determined by plaque assay and further passaged in Calu-3 or Vero cells at an MOI of 0.01 in DMEM supplemented with 10% FBS. P1 viruses were harvested at 48 h post-infection by freezing at −80°C. SARS-CoV-2 BAC with S813K or S813V mutations were recovered as described above.

For the antibody resistance selection, bFP antibodies were incubated with the P0 viruses at a concentration that corresponds to PRNT_90_ at 37°C for 1 h. The amount of P0 viruses used corresponds to the amount needed for infection at an MOI of 0.01 in a T75 flask. Calu-3 or Vero cells were then infected with the virus inoculum for 1 h in the presence of 230 μg/mL COV44-62 antibody or 330 μg/mL COV44-79 antibody. The virus inoculum was removed after virus adsorption and cells were washed with PBS before supplementing culture medium with 230 μg/mL COV44-62 antibody or 330 μg/mL COV44-79 antibody. Supernatant and cells were harvested at 48 h post-infection by freezing at −80°C.

### Sequencing library preparation

Viruses from different passages were inactivated in TRIzol (Thermo Fisher Scientific, catalog no. 15596026) for RNA isolation as specified by manufacturer’s protocol. Isolated RNA was subject to DNase I treatment (Thermo Fisher Scientific, catalog no. 18068015) and reverse-transcribed using the SuperScript IV First-Strand Synthesis System with random hexamers (Thermo Fisher Scientific, catalog no. 18091050). Region corresponding to residues 805-864 was amplified from the cDNA (post-selection) or the BAC mutant library (input) using KOD Hot Start DNA polymerase (MilliporeSigma, catalog no. 710863) per the manufacturer’s instruction with the following two primers: 5’-CAC TCT TTC CCT ACA CGA CGC TCT TCC GAT CT<u>T TTG GTG GTT TTA ATT TTT CAC AA</u>-3’ and 5’-GAC TGG AGT TCA GAC GTG TGC TCT TCC GAT CT<u>TTGA GCA ATC ATT TCA TCT GTG AG</u>-3’. Sequences complementary to the cDNA are underlined, whereas the rest of the sequences correspond to Illumina adapter sequence. An additional PCR was performed to add the rest of the Illumina adapter sequence and index to the amplicon using primers: 5’-AAT GAT ACG GCG ACC ACC GAG ATC TAC ACX XXX XXX XAC ACT CTT TCC CTA CAC GAC GCT-3’ and 5’-CAA GCA GAA GAC GGC ATA CGA GAT XXX XXX XXG TGA CTG GAG TTC AGA CGT GTG CT-3’. Positions annotated by an X represent the nucleotides for the index sequence. The final PCR products were purified by PureLink PCR purification kit (Thermo Fisher Scientific, catalog no. K310002) and submitted for next-generation sequencing using Illumina MiSeq PE250.

### Sequencing data analysis

Next-generation sequencing data were obtained in FASTQ format. Forward and reverse reads of each paired-end read were merged by PEAR^64^. The merged reads were parsed by SeqIO module in BioPython^65^. Primer sequences were trimmed from the merged reads. Trimmed reads with lengths inconsistent with the expected length were discarded. The trimmed reads were then translated to amino acid sequences, with sequencing error correction performed at the same time as previously described^61^. Amino acid mutations were called by comparing the translated reads to the WT amino acid sequence. Frequency (F) of a mutant i within sample s of replicate k was computed for each replicate as follows:

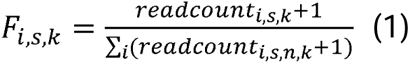

Mutants with a frequency of <0.01% in the BAC mutant library were discarded.

Enrichment score (ES) of a mutant i in replicate k was calculated as follows:

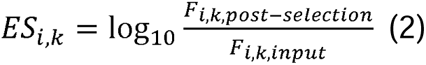

Fitness value (W) of a mutant i in replicate k was calculated as follows:

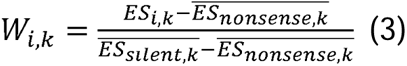

where 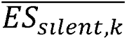 and 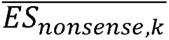 represent the average ES for silent and nonsense mutations, respectively, in replicate k.

The final fitness value for each mutant was the average W of the two replicates. The mutational tolerance for each residue was computed as the average fitness value of mutations at the given residue.

### Antibody expression and purification

The heavy chain and light chain of the indicated antibodies were cloned into phCMV3 plasmids in an IgG1 or Fab format with a mouse immunoglobulin kappa signal peptide. Plasmids encoding the heavy chain and light chain of antibodies were transfected into Expi293F cells using an Expifectamine 293 transfection kit (Gibco) in a 2:1 mass ratio following the manufacturer’s protocol. Supernatant was harvested 6 days post-transfection and centrifuged at 4000 × *g* for 30 min at 4°C to remove cells and debris. The supernatant was subsequently clarified using a polyethersulfone membrane filter with a 0.22 μm pore size (Millipore).

CaptureSelect CH1-XL beads (Thermo Scientific) were washed with MilliQ H_2_O thrice and resuspended in 1× PBS. The clarified supernatant was incubated with washed beads overnight at 4°C with gentle rocking. Then, flowthrough was collected, and beads washed once with 1× PBS. Beads were incubated in 60 mM sodium acetate, pH 3.7 for 10 min at 4°C. The eluate containing antibody was buffer-exchanged into 1× PBS and further purified by size-exclusion chromatography using Superdex 200 XK 16/100 column in 1× PBS. Antibodies were stored at 4°C.

### Biolayer interferometry binding assay

Binding assays were performed by biolayer interferometry (BLI) using an Octet Red instrument (FortéBio). Briefly, an N-terminally biotinylated peptide of SARS-CoV-2 S (808-DPSKPSKRSFIEDLLFNKVT-827) as well as a version with F823Y mutation at 50 μg/ml in 1x kinetics buffer (1x PBS, pH 7.4, 0.01% BSA and 0.002% Tween 20) were loaded onto SA biosensors and incubated with the COV44-62 and COV44-79 Fabs at 33.3 nM, 100 nM, and 300 nM. The assay consisted of five steps: 1) baseline: 60 s with 1x kinetics buffer; 2) loading: 180 s with biotinylated peptides, 3) baseline: 60 s with 1x kinetics buffer; 4) association: 180 s with Fabs; and 6) dissociation: 180 s with 1x kinetics buffer. For estimating the exact KD, a 1:1 binding model was used.

### Pseudovirus virus entry assay

Full-length SARS-CoV-2 S gene (GenBank: NC_045512.2) was synthesized by Genscript. as human codon-optimized cDNAs, and inserted into pcDNA3.1 expression vector^66^. C9-tagged versions of the S genes were generated by replacing the 3’-terminal 19 codons with linker and C9 codons (GSSGGSSG-GGTETSQVAPA)^67^. All S recombinants were constructed via gene fragment Assembly (New England Biolabs, catalog #: E2621S).

pHEF-VSVG-Indiana was constructed previously^68^. VSVGΔG-fluc-G pseudoviral particles (VSVpps^69^) stock was made as previously described^70^. Briefly, HEK293T cells were transfected with VSV-G. Next day, seed VSVΔG-G particles were inoculated onto the transfected cells for 2 h. The cells were rinsed three times with FBS-free DMEM medium and replenished with fresh media. After a 48-h incubation period, media were collected and clarified (300 × g, 4°C, 10 min then 3000 × g, 4°C, 10 min). To obtain purified viral particles, clarified VLP-containing media were laid on top of a 20% w/w sucrose cushions and viral particles were purified via slow-speed pelleting (SW28, 6500 rpm, 4°C, 24 h). The resulting pellet was resuspended in FBS-free DMEM to 1/100 of the original volumes. Concentrated particle stocks were stored at −80°C until used.

VSVpps bearing various recombinant SARS-CoV-2 S proteins were used to infect different cell types. For protease/antibody inhibition experiments, cells were pre-incubated with serial dilutions of camostat, E64D, or antibodies for 1 h at 37°C before VSVpp inoculation. Inoculation was allowed to infect cells for 2 h, then cells were rinsed 3 times and replenished with cell culture media (with 10% FBS). Following overnight incubation, cells were lysed by lysis buffer (25 mM Tris-phosphate pH 7.8, 2 mM dithiothreitol, 2 mM 1,2-diaminocyclohexane-*N*,*N*,*N*′-tetraacetic acid, 10% glycerol, 1% Triton X-100). Firefly luciferase (VSVpp) activity was recorded by a Veritas microplate luminometer after addition of substrate (1 mM d-luciferin, 3LmM ATP, 15LmM MgSO_4_·H_2_O, 30 mM HEPES pH 7.8).

### Western blot analysis

Samples in SDS solubilizer (0.0625 M Tris·HCl pH 6.8, 10% glycerol, 0.01% bromophenol blue, 2% SDS, and 2% 2-mercaptoethanol) were heated at 95°C for 5 min, electrophoresed through 8% polyacrylamide-SDS gels, transferred to nitrocellulose membranes (Bio-Rad), and incubated with rabbit polyclonal anti-SARS-CoV-2-S1 (SinoBiological, catalog #: 40591-T62), mouse anti-C9 (EMD Millipore, catalog #: MAB5356), mouse monoclonal anti-VSV-M (KeraFast, catalog #: EB0011). After incubation with appropriate HRP-tagged secondary antibodies and chemiluminescent substrate (Thermo Fisher), or purified LgBiT-substrate cocktail (Promega), the blots were imaged and processed with a FluorChem E (Protein Simple).

### Structural modelling

FoldX^52^ was used to model the structural and protein stability effects of mutation F823Y. The published structures of SARS-CoV-2 bFP in complex COV44-62 (PDB 8D36)^17^ and COV44-79 (PDB 8DAO)^17^ were used as input.

### Sequence alignment

Sequence alignment was performed using (http://www.bioinformatics.org/sms/multi_align.html)^71^. Sequences were downloaded from NCBI GenBank database (www.ncbi.nlm.nih.gov/genbank)^72^. Genbank IDs for the S sequences used are as follows:

ABB90529.1: Human coronavirus 229E (HCoV-229E)

YP_003767.1: Human coronavirus NL63 (HCoV-NL63)

ADN03339.1: Human coronavirus HKU1 (HCoV-HKU1)

AIX10756.1: Human coronavirus OC43 (HCoV-OC43)

YP_001039971.1: Rousettus bat coronavirus HKU9 (BatCoV-HKU9)

ABF65836.1: Severe acute respiratory syndrome-related coronavirus (SARS-CoV)

QHD43416.1: Severe acute respiratory syndrome coronavirus 2 (SARS-CoV-2)

AHX00731.1: Middle East respiratory syndrome-related coronavirus (MERS-CoV)

YP_001876437.1: Beluga whale coronavirus SW1 (BWCoV-SW1)

AHB63508.1: Bottlenose dolphin coronavirus HKU22 (BDCoV-HKU22)

AFD29226.1: Night heron coronavirus HKU19 (NHCoV-HKU19)

AFD29187.1: Porcine coronavirus HKU15 (PDCoV-HKU15)

## Code availability

Custom python scripts for all analyses have been deposited to: https://github.com/nicwulab/SARS2_FP_DMS

## Data availability

Raw sequencing data have been submitted to the NIH Short Read Archive under accession number: BioProject PRJNA910585.

