## Supplementary data for "Functional and antigenic characterization of SARS-CoV-2 spike fusion peptide by deep mutational scanning"

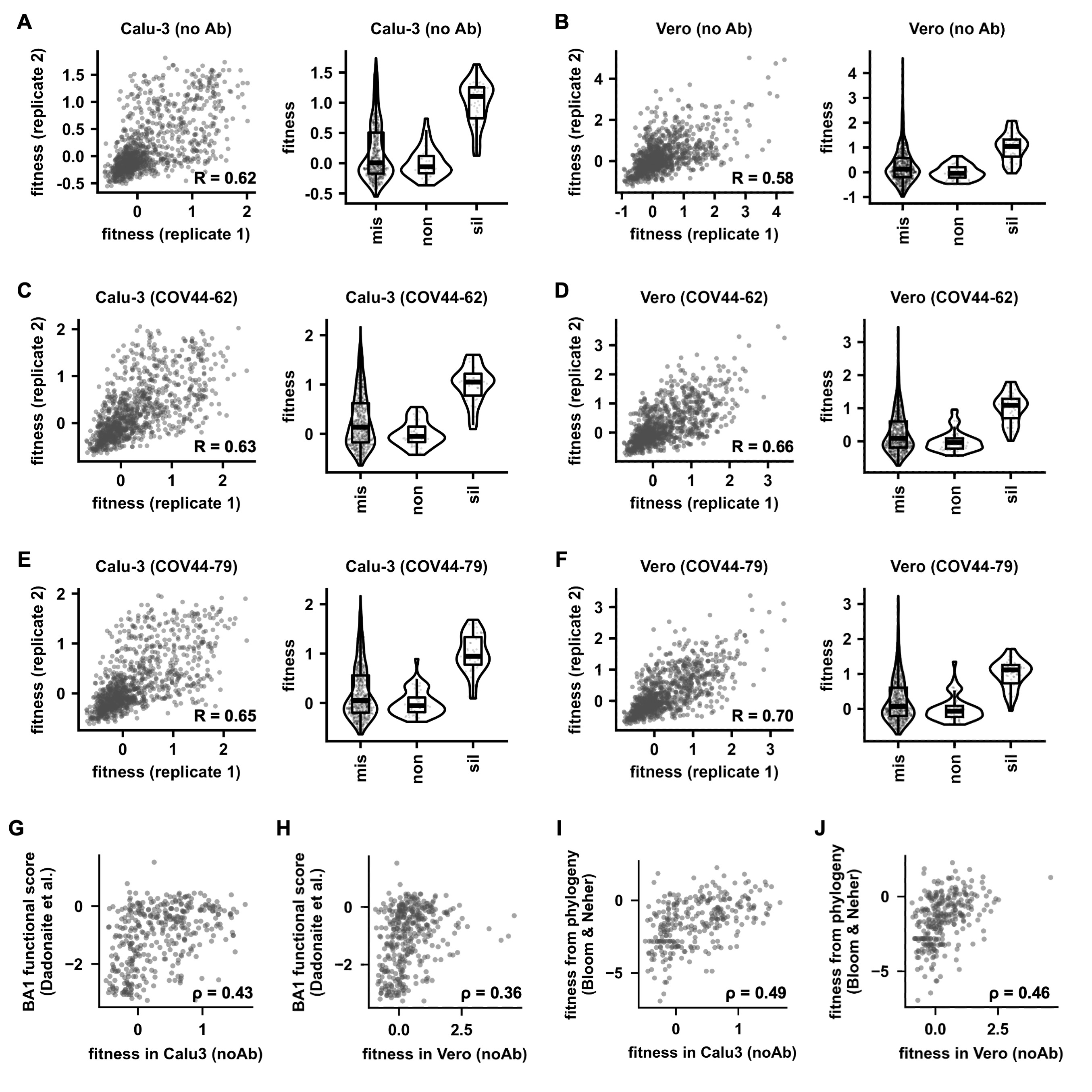

**Figure S1. Reproducibility and quality of the deep mutational scanning data.** Correlation of fitness values for individual mutations between two biological replicates is shown as a scatterplot (left panel). The distributions of fitness values for missense mutations (mis), nonsense mutations (non), and silent mutations (sil) are shown as a stripchart overlayed with a violin plot (right panel). Of note, silent mutations represent nucleotide variants that encode the WT amino-acid sequences but were different from the WT nucleotide sequence. Deep mutational scanning was performed in six conditions, namely **(A)** Calu-3 cells with no antibody selection, **(B)** Vero cells with no antibody selection, **(C)** Calu-3 cells with COV44-62 antibody selection, **(D)** Vero cells with COV44-62 antibody selection, **(E)** Calu-3 cells with COV44-79 antibody selection, and **(F)** Vero cells with COV44-79 antibody selection. **(G-H)** Previously, Dadonaite et al. measured the effects of mutations in BA.1 S on virus entry [S1]. Correlations between the functional scores reported by Dadonaite et al. [S1] and **(G)** fitness values in Calu-3 cells, or **(H)** Vero cells are shown. **(I-J)** Previously, Bloom and Neher computed the fitness effects of mutations to all SARS-CoV-2 proteins using a phylogeny-based approach [S2]. Correlations between the fitness computed by Bloom and Neher [S2] and **(I)** fitness values in Calu-3 cells, or **(J)** Vero cells are shown. **(G-I)** Only missense mutations were analyzed. Spearman's rank correlation coefficients (ρ) are indicated.

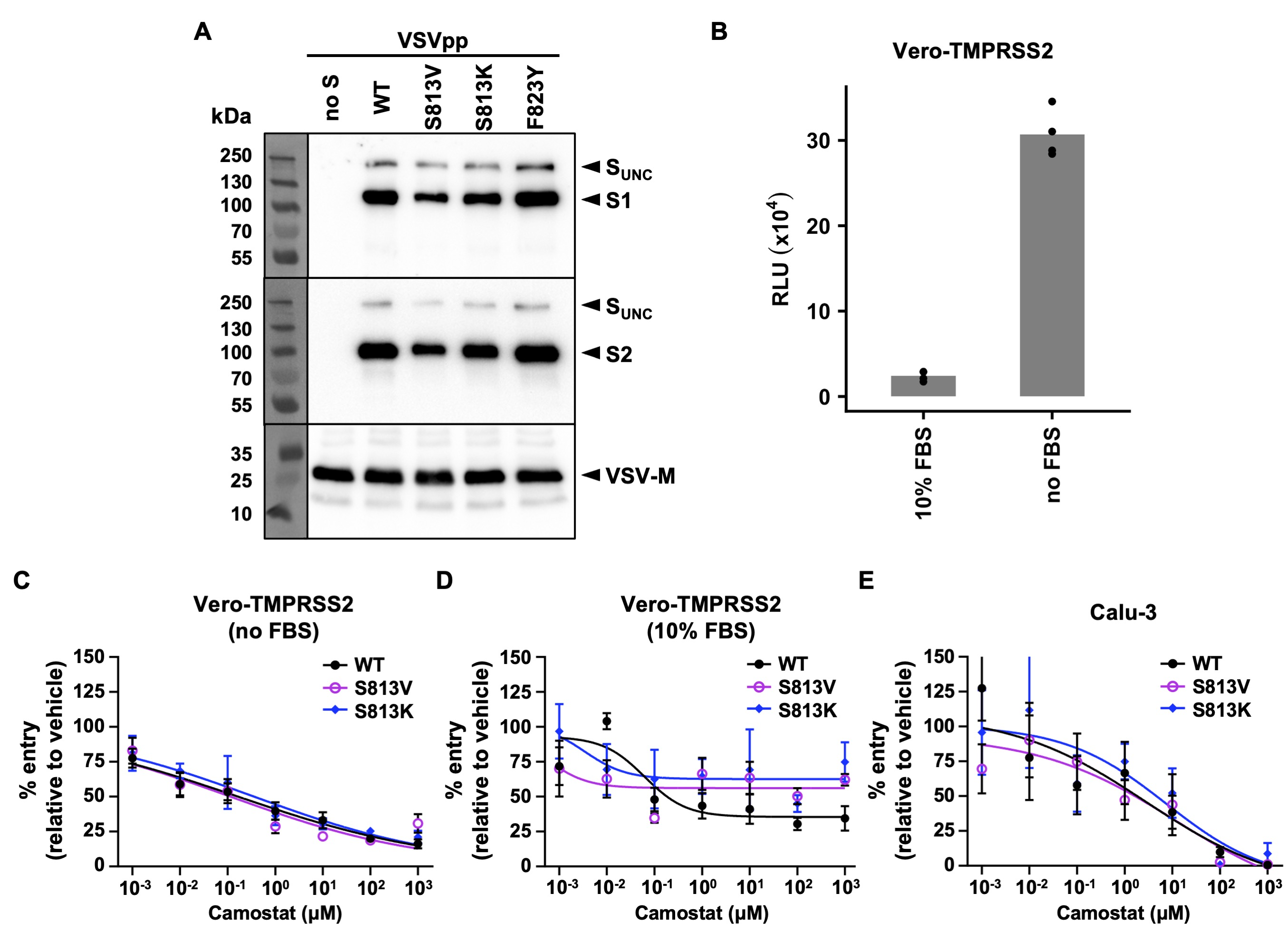

**Figure S2. Characterizing SARS-CoV-2 S mutations using VSVpps. (A)** Western blot analysis of VSVpps bearing various S constructs. Uncleaved S (S_UNC_), S1, S2, and VSV-M are labeled. **(B)** Vero-TMPRSS2 cell entry in the absence or presence of FBS by VSVpps bearing SARS-CoV-2 S (WT) was measured by the relative light unit (RLU) in a luciferase assay. Each bar represents the mean of four biological replicates. Each datapoint represents one biological replicate. **(C and D)** The effects of camostat on Vero-TMPRSS2 cell entry of VSVpps bearing various SARS-CoV-2 S constructs, **(C)** in the absence or **(D)** presence of FBS. **(E)** The effects of camostat on Calu-3 cell entry of VSVpps bearing various SARS-CoV-2 S constructs. Mean and SEM of four biological replicates are depicted.

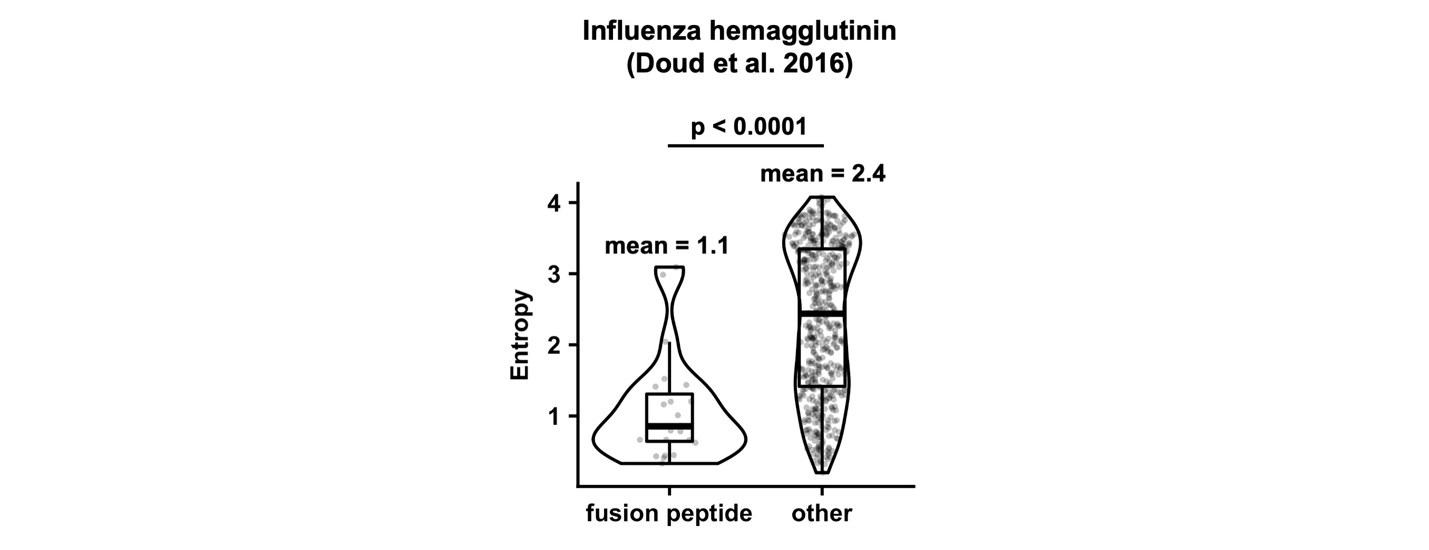

**Figure S3. Mutational tolerance of influenza hemagglutinin fusion peptide.** A previous deep mutational scanning study reported the mutational tolerance of each amino acid residue in the influenza H1N1 hemagglutinin [S3]. This previous study quantified mutational tolerance of each amino acid residue as entropy. A lower entropy value indicates lower mutational tolerance. The distributions of entropy for the 23 residues in the fusion peptide of influenza hemagglutinin and non-fusion peptide residues are shown as a stripchart overlayed with a violin plot. The indicated p-value is computed by Wilcoxon rank-sum test.

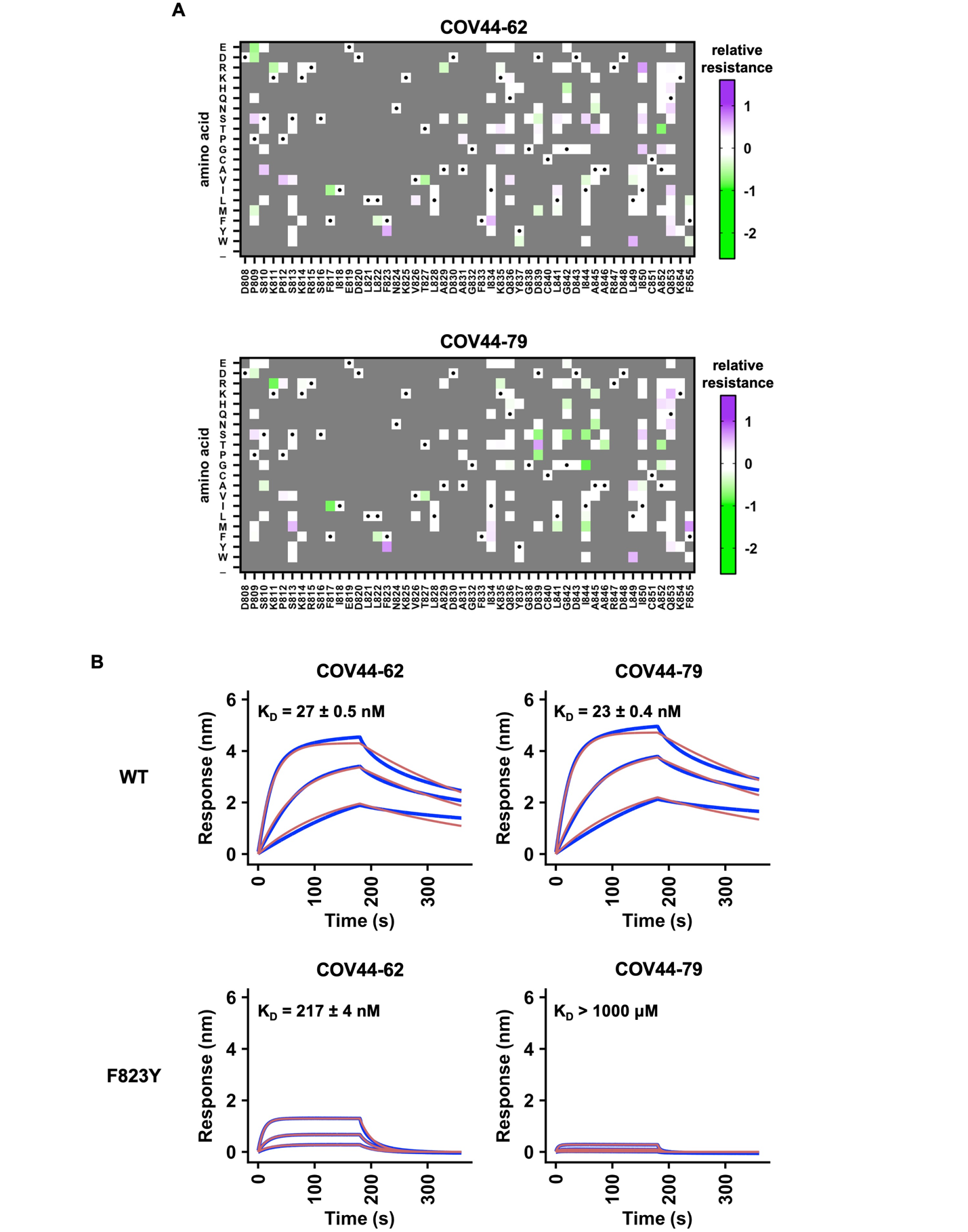
**Figure S4. F823Y mutation weakens the binding of bFP antibodies. (A)** Relative resistance for each mutation against 230 μg/mL COV44-62 antibody or 330 μg/mL COV44-79 antibody in Calu-3 cells is shown as heatmaps. Relative resistance for WT is set as 0. Mutations with a fitness value of less than 0.75 are shown as gray. Amino acids corresponding to the WT sequence are indicated by the black dots. **(B)** Binding kinetics of COV44-62 Fab or COV44-79 Fab against WT or F823Y peptide that contained residues 808 to 827 were measured by biolayer interferometry (BLI). Y-axis represents the response. Blue lines represent the response curve and red lines represent the 1:1 binding model. Binding kinetics were measured for three concentrations of Fab at 3-fold dilution ranging from 300 nM to 33.3 nM.

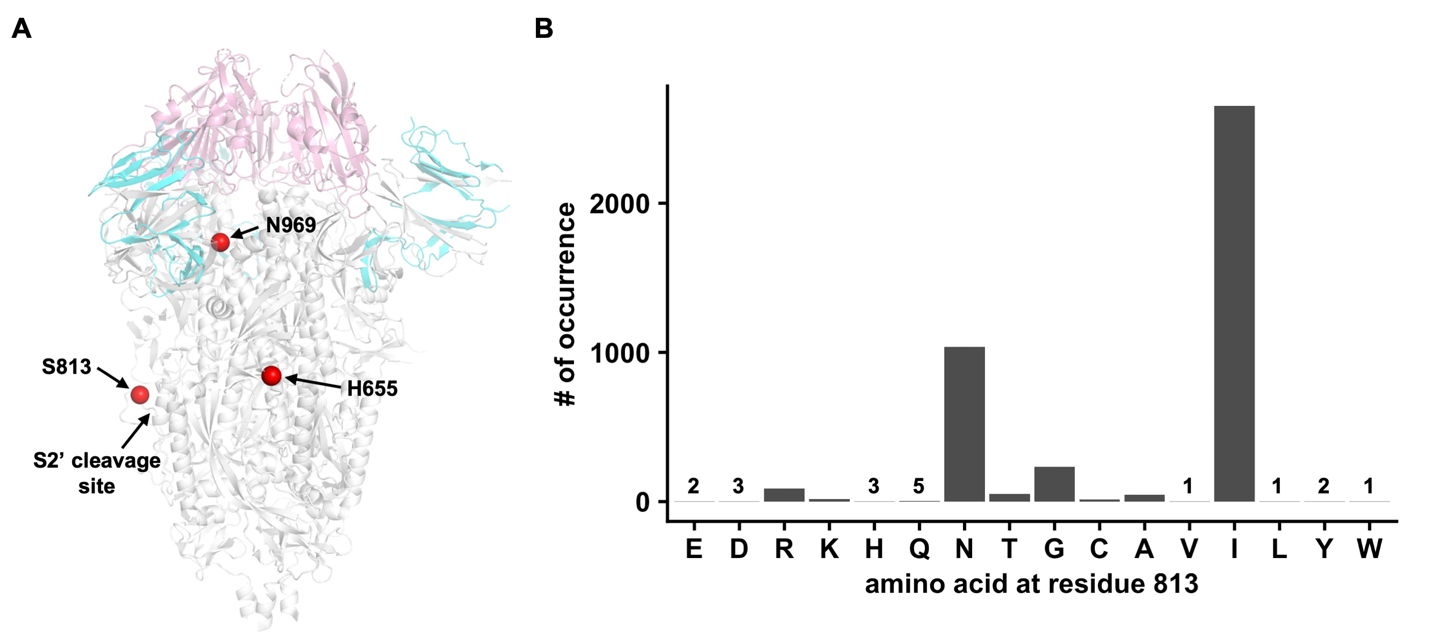
**Figure S5. Frequency of natural mutations at residue 813. (A)** The Cαs of residues 655, 813 and 969 are shown in red spheres on the SARS-CoV-2 spike structure (PDB 6VXX) [S4]. **(B)** Occurrences of different amino acid mutations at residue 813 among 15 million SARS-CoV-2 genomes on GISAID are shown. The wild-type variant Ser (S) is not shown. Occurrence of less than 10 is indicated.

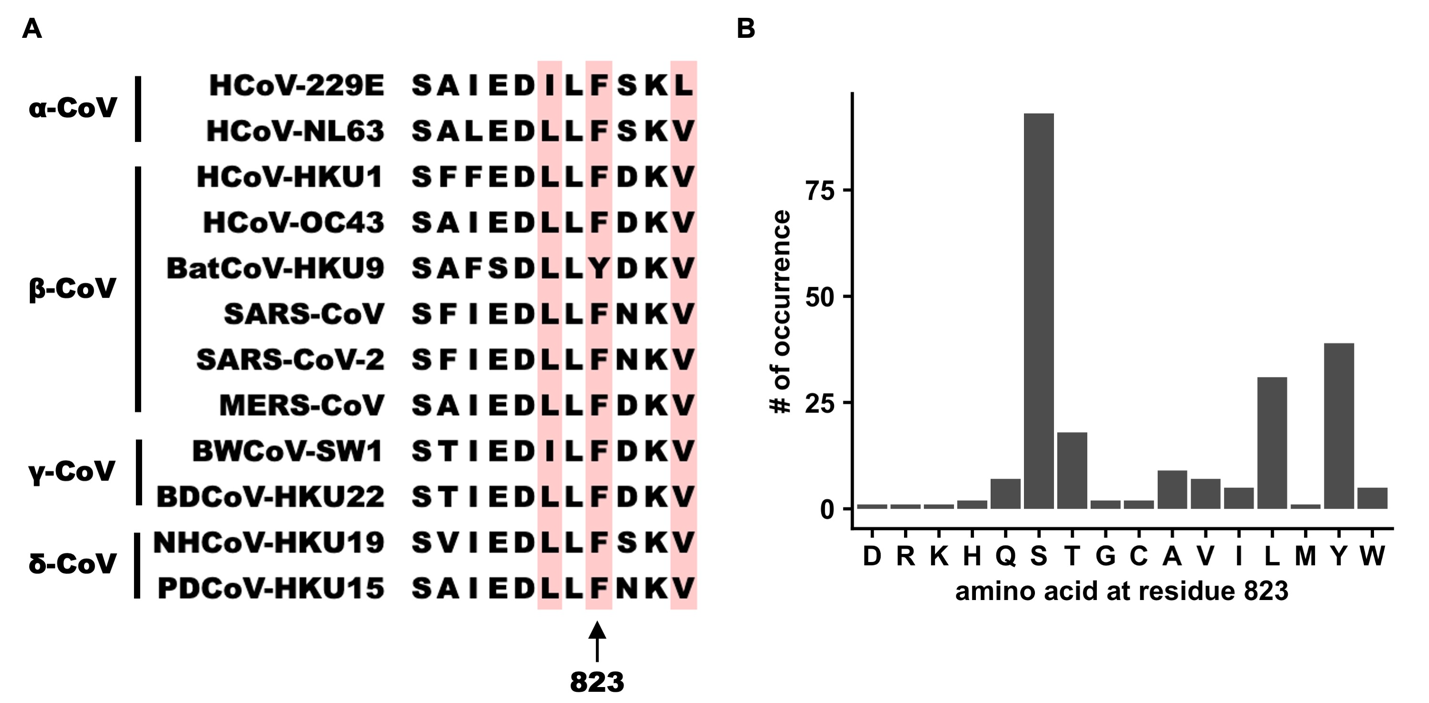

**Figure S6. Natural occurrence of F823Y. (A)** Multiple sequence alignment of the first 11 residues of bFP from different strains that represent four coronavirus subgroups (α, β, γ, and δ). Residues that are not completely conserved among these sequences are highlighted in pink. **(B)** Occurrences of different amino acid mutations at residue 823 among 15 million SARS-CoV-2 genomes on GISAID are shown. The wild-type variant Phe (F) is not shown.
